# Neutrophils Mediate Kidney Inflammation Following Acute Skin Exposure to UVB Light

**DOI:** 10.1101/2020.05.25.115204

**Authors:** Sladjana Skopelja-Gardner, Joyce Tai, Xizhang Sun, Lena Tanaka, James A. Kuchenbecker, Jessica M. Snyder, Paul Kubes, Tomas Mustelin, Keith B. Elkon

**Affiliations:** Division of Rheumatology, University of Washington, Seattle, WA, USA; Department of Ophthalmology, University of Washington, Seattle, WA, USA; Department of Comparative Medicine, University of Washington, Seattle, WA, USA; Calvin, Phoebe and Joan Snyder Institute for Chronic Diseases, University of Calgary, Alberta, CA; Department of Immunology, University of Washington, Seattle, WA, USA; Center for Innate Immunity and Immune Disease, University of Washington, Seattle, WA, USA

## Abstract

Photosensitivity to ultraviolet (UV) light affects up to ~80% of lupus patients and can exacerbate local skin disease as well as systemic disease, including lupus nephritis. While neutrophils have been implicated in local tissue injury in lupus in response to immune complex deposition, whether and how they play a role in photosensitivity induced systemic disease is unknown. Here, we show that following skin exposure to UV light, neutrophils migrate not only to the skin, but also to the kidney, in an IL-17A-dependent manner. Kidney infiltrating neutrophils produced reactive oxygen species and their presence was associated with upregulation of endothelial adhesion molecules and inflammatory cytokines as well as the induction of kidney injury markers, including transient proteinuria. Neutrophils were responsible for inflammation and renal injury as demonstrated by experiments that inhibited neutrophil mobilization. Exploiting a mouse model containing photoactivatable immune cells, we observed that a subset of neutrophils found in the kidney had transited through UV light-exposed skin suggesting reverse transmigration. These findings demonstrate that neutrophils mediate transient kidney injury following skin exposure to UV light and, coupled with observations identifying similar neutrophil phenotypes in human lupus, could provide a mechanistic link to explain sun-induced systemic lupus flares.

## Introduction

Sensitivity to ultraviolet (UV) rays from the sun is a well-recognized feature of systemic lupus erythematosus (SLE)^1-3^. Exposure to UV light triggers both local inflammation in the skin and has been associated with systemic disease flares, including lupus nephritis (LN), in SLE patients^4-6^. How photosensitivity in the skin leads to systemic manifestations remains poorly understood. Shared gene signatures in the skin and kidney of SLE patients^7,8^ suggest a common pathogenesis. We recently observed that acute skin exposure to UV light triggers both a local and a systemic type I interferon (IFN-I) response^9^. However, the effects of skin exposure to UV light on kidney function were not examined.

Recent findings indicate that neutrophils play an important role in both local and systemic lupus disease. Neutrophils are present in the skin lesions of SLE patients^10-12^ as well as in the kidney tissue of patients with LN^13,14^. High expression of a neutrophil gene signature is a strong predictor of active SLE, including nephritis and cutaneous flares^15-17^. A specific subset of pro-inflammatory low-density granulocytes (LDGs) are increased in SLE, particularly in patients with skin disease^18^. Neutrophils can contribute to tissue injury, for example by releasing proteases, reactive oxygen species (ROS), or via extrusion of neutrophil extracellular traps (NETs)^14,19-21^. Since UV light is known to induce rapid neutrophil infiltration into the skin,^22-24^ we investigated how inflammation triggered by skin exposure to UV light influenced neutrophil migration and function both at the local site of injury and in remote locations, particularly the kidney.

## Results

### UV light-induced sterile inflammation in the skin is associated with neutrophilia and neutrophil entry into organs

To investigate whether neutrophils participate in the systemic inflammatory response to UVB light (a single dose of 500mJ/cm^2^), we examined the kinetics of neutrophil accumulation in UVB irradiated skin as well as in spleen, kidney, and lung of C57BL/6 J (B6) mice, a strain that best mimics cutaneous changes to UVB light in human skin (see **Fig. 1A**)^25^. Lung was chosen as a control organ as no association between photosensitivity and clinical lung disease have been reported in SLE. 500 mJ/cm^2^ of UVB light in B6 mice has been defined as 2 minimal erythematous doses,^26,27^ the reference dose used for human phototesting^28^. Following skin exposure to UVB light, neutrophil numbers in the skin increased 7-fold within 24 h and remained elevated for 6 days, compared to baseline pre-UV levels (D-1) (**Fig. 1B**). This was accompanied by neutrophil egress from the bone marrow and a 5-fold increase in circulating blood neutrophils on days 1-6 (**Fig. 1C, D**). Intriguingly, in the same time frame neutrophils increased in the spleen, lung, and kidney (**Fig. 1E-G**, Supplementary Fig. 1A). The precise kinetics varied somewhat between organs, reaching peak on day 1 in the lung and spleen and on days 2-6 in the kidney (**Fig. 1E-G**). In the kidneys, neutrophils mostly localized to the tubulointerstitial area, including the vasculature, as indicated by neutrophil elastase (NE) staining in proximity to, or co-localized with, platelet/endothelial cell adhesion molecule-1 (PECAM-1/CD31) (**Fig. 1H**). Representative images in Fig. 1H show neutrophils in or around the interstitial vasculature (full white arrows) and in the interstitium (dotted white arrows), with occasional neutrophils also found in the glomeruli (yellow arrows). Similar localization was detected with ant-Ly6G IF staining (Supplementary Fig. 1B).

**Figure 1:**
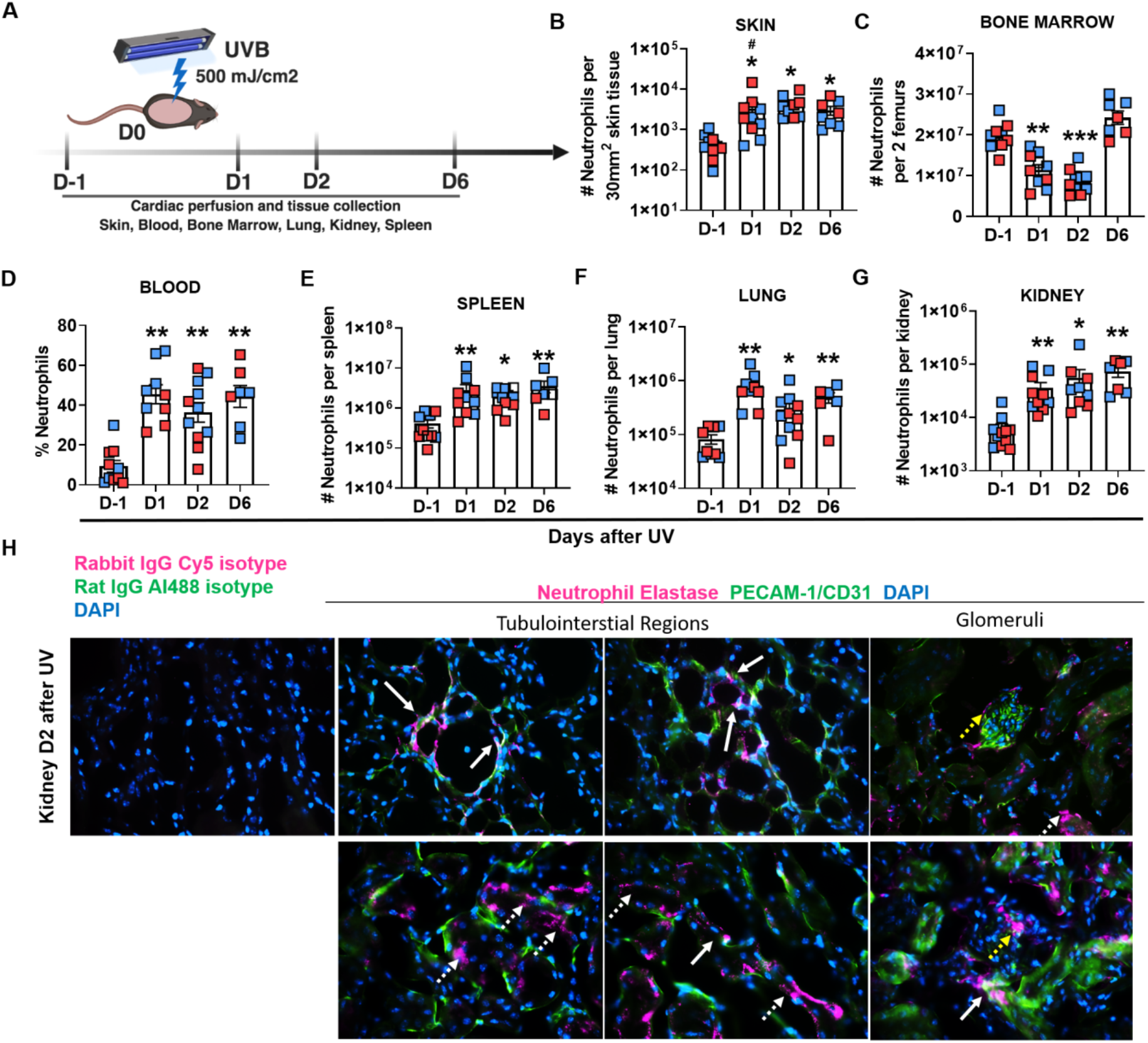
Skin exposure to UVB light triggers a neutrophilia, neutrophil infiltration of skin and peripheral organs including the kidney. **(A)** Following a single exposure of skin to 500 mJ/cm^2^ UVB light, neutrophils were quantified in the blood, skin, lung, spleen, and kidney at the time points shown. Following whole body perfusion, cells were isolated from the sites shown in **(B-G)** and the number of neutrophils (Ly6C+Ly6G+ cells) evaluated in the CD45+ live cells gate in female (red) and male (blue) mice on days (D) 1, 2, and 6 after UV exposure. Differences in neutrophil numbers on different days after exposure to UVB light were determined relative to non-irradiated controls (D-1) using Student’s t-test (n=6 −10 per group; *p<0.05, **p<0.01, ***p<0.001; #p< 0.05 comparing fold increase in male vs. female mice). Representative flow cytometry plots for each organ are shown in fig.S5. (**H**) Immunofluorescence staining of frozen kidney sections on day 2 (D2) after UV exposure demonstrating neutrophil elastase (Cy5, magenta), PECAM-1/CD31 (Al488, green), and nuclear DAPI (blue) staining in the tubulointerstitium and glomeruli. Neutrophil elastase (NE) staining in vascular endothelium is denoted by white full arrow (colocalization of NE and CD31, yellow); NE staining in the interstitial tissue denoted by dotted white arrows; NE staining in glomeruli denoted by yellow arrows. Isotype control staining is shown in the leftmost panel. Magnification = 40X.

To determine whether other innate immune cells showed local and systemic responses similar to neutrophils, we examined the distribution of monocytes (CD11b+Ly6C+Ly6G-) at the same time points after UVB light exposure. While monocytes were also recruited from the blood and bone marrow pools into the skin, little or no monocyte migration to the lung or spleen was observed and only small numbers of infiltrating monocytes were detected in the kidney on day 6 after UVB exposure (~2-fold vs. ~10.5-fold increase in kidney neutrophils) (Fig. 1G, Supplementary Fig. 2). No increase in monocyte numbers was detected in the lung or spleen (Supplementary Fig. 2), suggesting that the systemic response to UV injury was relatively selective for neutrophils.

Infiltration of neutrophils into the skin was accompanied by histologic changes including early inflammation of the dermis and epidermal cell death (days 1-2), followed by development of acanthosis and hyperkeratosis, with serocellular crust formation in some cases (day 6) (Supplementary Fig. 3). Notably, no open skin lesions appeared after exposure to UVB light, suggesting that the observed immune cell infiltration occurred under sterile conditions, although a contribution by an altered skin microbiome cannot be excluded^26^.

Together, these findings demonstrate that exposure of the skin to a single dose of UVB light mobilizes neutrophils to both the local site of inflammation as well as to internal organs, accompanied by blood neutrophilia. While overall migratory patterns were similar in both male and female mice, neutrophil infiltration into the skin was more rapid in females compared to males (Fig. 1B). In contrast, epidermal injury by tape stripping did not lead to neutrophil migration to the kidney despite their infiltration into the injured skin (Supplementary Fig. 4), indicating that systemic neutrophil dissemination is not common to all forms of interface dermatitis.

### IL-17-A promotes neutrophil recruitment following skin exposure to UV light

A survey of chemotactic and inflammatory mediators in the skin revealed significant upregulation in the gene expression of neutrophil chemokines and inflammatory cytokines *cxcl1, cxcl5/6*, as well as *g-csf*, and *il-lβ* 1-2 days after UV exposure (**Fig. 2A-D**). The rapid and persistent increase in expression of *s100A9* matched the kinetics of neutrophil infiltration into the skin (Fig. 1B, Supplementary Fig. 5) whereas the cytokines *il-6, tnf*, and *il-33* were transiently increased and back to baseline expression by day 6 (Supplementary Fig. 5). In contrast, *il-36a* was elevated only on day 6, perhaps reflecting a reparative function^29,30^. The transient (1-2 days) presence of monocytes in the skin (Supplementary Fig. 5) paralleled the upregulation of monocyte-specific chemokines *ccl4* and *ccl2* (Supplementary Fig. 5). Similar to the accumulation of neutrophils in the skin of female mice (Fig. 1B), monocyte accumulation in the skin was also faster in females than males (Supplementary Fig. 2).

**Figure 2:**
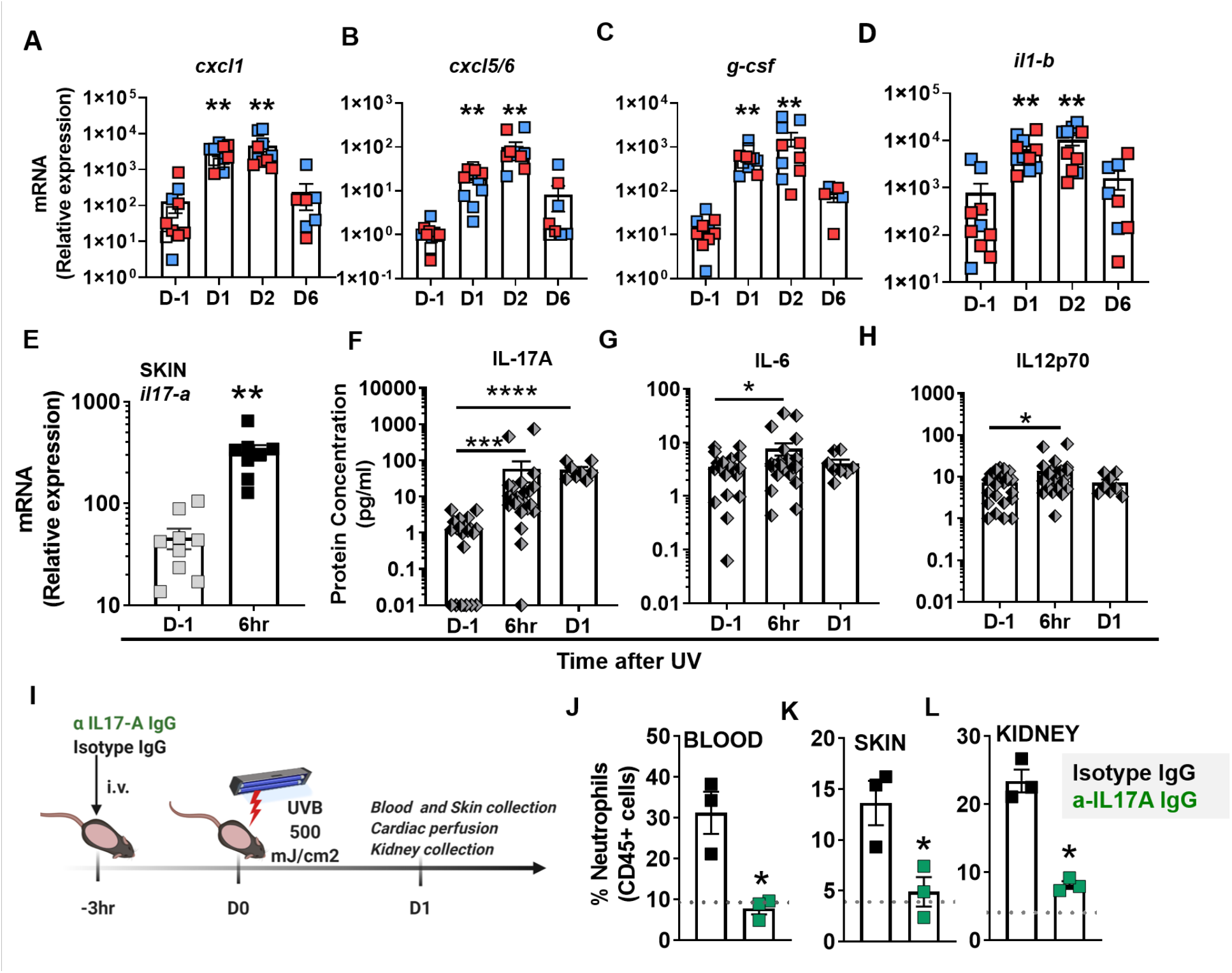
IL17-A mediates local and systemic neutrophil recruitment after skin exposure to UVB light. **(A-D)** Relative mRNA expression in skin obtained from female (red) and male (blue) mice was quantified by QPCR using the primers listed in Table 1 and normalized to *18S* transcript levels. Bars represent mean relative expression + SEM for all samples combined. **(E-G)** Multiplex analysis of protein concentration in plasma samples collected 6 hr or one day (D1) after UV (E) IL17-A, (F) IL-6, and (G) IL-12p70. (**H**) Il17A gene expression in the skin 6hr after UV exposure was quantified as in (A-D). Significant changes in gene expression and plasma protein levels at different time-points after exposure to UV light were determined relative to those of non-irradiated controls (D-1) using Student’s t-test (A-D, H, n=8-10 and E-G, n=8-25 mice per group; *p<0.05, **p<0.01, ***p<0.001). (**I**) Shaved B6 mice were treated i.v. with 100 μg anti-IL17A IgG (green) or isotype control IgG (black) 3hr before exposure to a single dose of UVB light (500 mJ/cm^2^). One day after UV exposure (D1), neutrophils were quantified in blood (**J**), skin (**K**), and kidney (**L**) by flow cytometry as in Fig. 1. Dotted grey lines represent average baseline neutrophil percentage in blood, skin, and kidney of non-injected mice. Statistical differences between treatment groups were determine by paired Student’s t-test (n = 3, *p < 0.05).

UV skin exposure was accompanied by rapid (6hr) 12-fold induction in *il-17a* gene expression in the skin (**Fig. 2E**), as well as a subsequent >1000-fold cutaneous induction in *g-csf* expression during days 1 and 2 (Fig. 2C). To determine whether this induction of these cytokines was relevant to neutrophil mobilization, we first quantified cytokine protein concentrations in the blood, which revealed a robust and sustained (10- to 100 fold) increase in the concentration of plasma IL-17A 6-24 hrs after exposure to UV light (**Fig. 2F**), together with a transient (peak at 6 hr) increase in IL-6 and IL-12, cytokines that may be downstream of IL-17A^31,32^ (**Fig. 2G, H**). No increase in the plasma concentrations of IFNg, GM-CSF, TNFa, IL-6, IL-10, IL-27, IL-23, or IL1a were observed (data not shown).

To determine if IL-17A is instrumental in UVB-induced neutrophil recruitment,^33,34^ we blocked the effect of IL-17A with a neutralizing antibody prior to UV exposure (**Fig. 2I**). Anti-IL-17A IgG significantly dampened blood neutrophilia (**Fig. 2J**) and attenuated neutrophil influx into both the exposed skin tissue (**Fig. 2K**) as well as the kidney (**Fig. 2L**). These findings demonstrate that local and systemic neutrophil recruitment in response to UVB light is mediated, at least in part, by IL-17A.

### Skin exposure to UV light causes rapid but reversible kidney injury

Next, we examined whether skin exposure to UVB light led to changes in the kidney (**Fig. 3A**). Since neutrophil influx into the skin of female B6 mice was faster and more striking than in male mice, and we previously showed a stronger inflammatory response to UV light in females (Fig. 1B, Supplementary Fig. 2,^9^), all subsequent studies were performed in female mice. As shown in **Fig. 3B**, kidney infiltrating neutrophils one day after skin exposure to UVB light produced reactive oxygen species (ROS), indicating that they were activated, compared to neutrophils present in the kidneys before UVB exposure of the skin (D-1). Extracellular localization of neutrophil elastase seen in the interstitium (Fig. 1H, bottom middle panel) and in the glomerulus (Fig. 1H, bottom right panel) is suggestive of neutrophil degranulation or NETosis, akin to that previously reported in the kidneys of SLE patients.^14^

**Figure 3:**
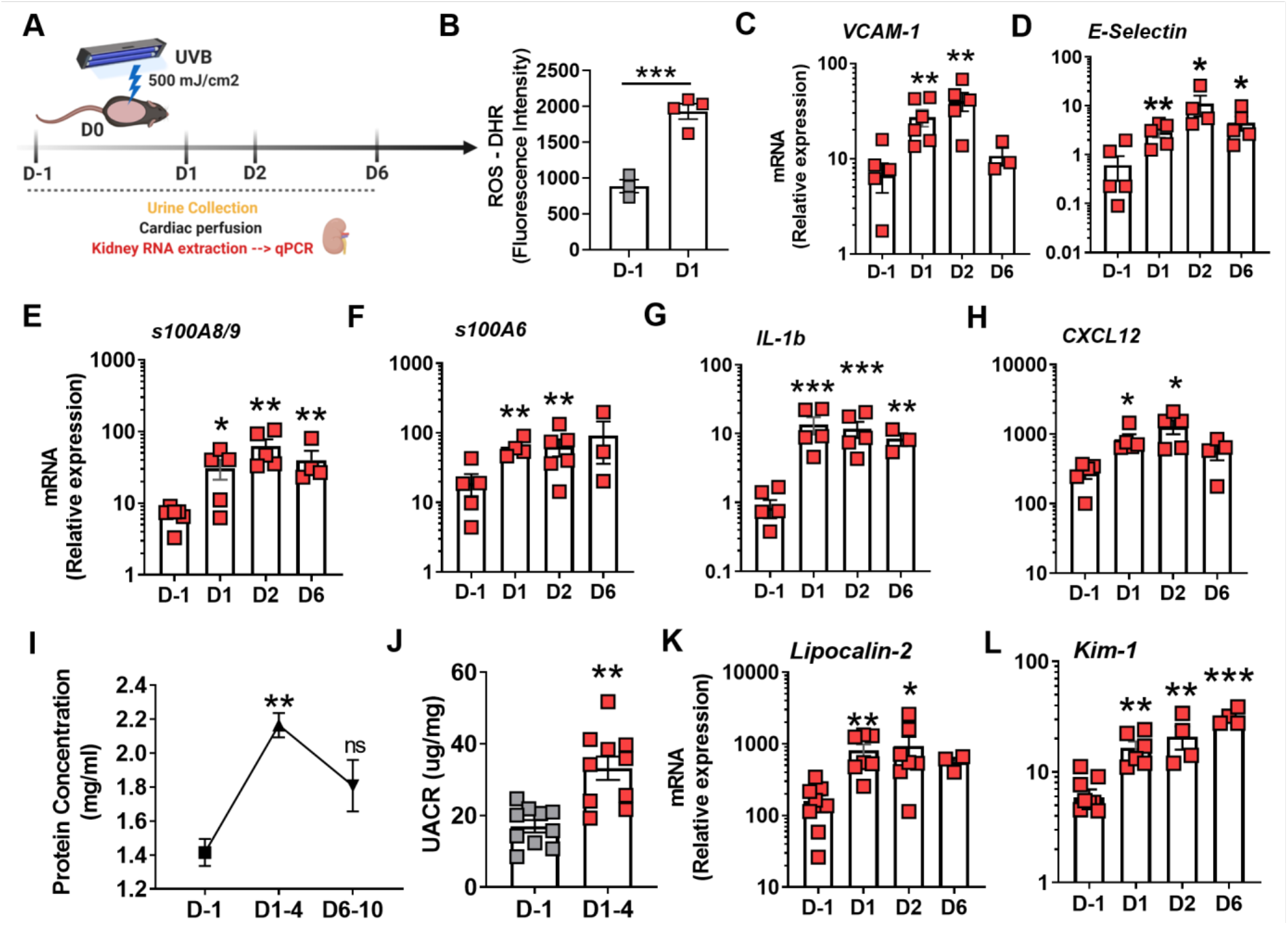
Skin exposure to UVB light triggers kidney inflammatory responses, transient proteinuria, and upregulation of kidney injury markers. **(A)** Following skin exposure to UVB light as in Fig.1, urine and kidney samples were collected at the time points shown. **(B)** Production of ROS by kidney infiltrating neutrophils 1day (D1) after skin exposure to UVB light was measured *ex vivo* by detection of 1,2,3 dyhidroxyrhodamine (DHR) fluorescence intensity and compared to cells from unexposed mice (D-1)**. (CD)** Gene expression of adhesion molecules *vcam-1* and *e-selectin*, (**E-H**) inflammatory *mediators s100a9, s100a6, il1b*, and *cxcl12* on different days following exposure to UVB light relative to the non-UV irradiated controls (D-1) was determined by QPCR. **(I-J)** Proteinuria was quantified by (I) Bradford assay and (J) Urine albumin/creatinine ratio (UACR) at the times shown after exposure to UVB light. Significant differences in proteinuria/UACR were determined relative to measurements from the urine of nonirradiated controls (D-1) using Student’s t-test (n=10 per time frame; **p<0.01). **(K-L)** Gene expression of renal endothelial injury markers (K) Lipocalin-2 and (L) Kim-1 in perfused kidney tissues over time following skin exposure to UV light was compared to non-UV exposed controls (D-1). Significance was determined by Student’s t-test (n=3-6 per group; *p<0.05, **p<0.01, ***p<0.001).

Gene expression analyses of perfused kidney tissues (Fig. 3A) revealed upregulation in the expression of the adhesion molecules *vcam-1* and *e-selectin* on days 1-2 after UVB (**Fig. 3C, D**). In contrast, *vcam-1* and *e-selectin* expression demonstrated a downward trend in the lung (Supplementary Fig. 6). In addition, we observed a >10-fold renal induction in *s100A9*, a neutrophilic mediator associated with kidney damage^35^, as well as increased *s100A6* expression, a calcium-binding protein associated with tubular injury^36^. (**Fig. 3E, F**). While *s100A9* induction was detected in the lung, likely indicative of neutrophil margination^37^, *s100A6* gene expression remained unchanged in the lung (Supplementary Fig. 6). The inflammatory response in the kidney was also associated with an early increase in *il-1β* expression (days 1 and 2 after UV), whereas there was minimal or no change in *tnf* or *il-6* (**Fig. 3G**, and data not shown). *Il-1β* expression was not detected in the lung until six days after skin exposure to UVB light (Supplementary Fig. 6). Rapid induction in *cxcl12* expression, a chemokine secreted by glomeruli and tubules and implicated in renal injury^38,39^, was also found in the kidney but not the lung (**Fig. 3H**, Supplementary Fig. 6). Therefore, UVB light-triggered skin injury instigates rapid neutrophil infiltration into the kidney associated with renal pro-inflammatory processes, while fewer changes are observed in the lung.

In addition to the pro-inflammatory neutrophil response observed in the kidney, both proteinuria and the urinary albumin/creatinine ratio increased in the first few days following skin exposure to UVB light (**Fig. 3I, J**). This rapid effect on kidney function was accompanied by upregulation in expression of *lipocalin-2* and kidney injury molecule 1 (*kim-1*) (**Fig. 3K, L**), markers of tubular kidney damage, also observed in lupus nephritis^40-42^. Despite the transient nature of proteinuria, expression of these kidney injury markers persisted up to day 6 (Fig. 3K, L), possibly reflecting tissue repair^43,44^.

### Neutrophils are responsible for kidney inflammation after skin exposure to UV light

To determine whether neutrophils were responsible for the inflammatory changes in the kidney following skin UVB light exposure, we inhibited neutrophil mobilization from the bone marrow by blocking granulocyte-colony stimulating factor (G-CSF) as outlined in **Fig. 4A**. Neutralizing G-CSF prevented blood neutrophilia as well as neutrophil migration to the kidney and skin following skin exposure to UVB light (**Fig. 4B, C** and not shown). Notably, as previously reported in different models^45-47^, blocking G-CSF did not alter monocyte numbers in the kidney (**Fig. 4D**). Decreased neutrophil migration to the kidney was accompanied by a significant reduction in the renal expression of the adhesion molecules *vcam-1* and *e-selectin* (**Fig. 4E, F**) as well as the inflammatory mediators *s100A9* and *il-1β* one day after UVB exposure (**Fig. 4G, H**). Furthermore, the tissue injury markers *lipocalin-2* and *kim-1* in the kidney were significantly decreased in UV light-exposed mice treated with G-CSF blocking antibody, relative to isotype treated, UV exposed animals (**Fig. 4I, J**). As we reported previously^9^, acute skin exposure to UVB light triggered an IFN-I response in the kidney (**Fig. 4K**). Blocking G-CSF significantly reduced, but did not completely abrogate, the IFN-score (Fig. 4K). Together, these data indicate that neutrophils are responsible for the inflammatory response and, directly or indirectly, contribute to type I interferon induction in the kidney following UV light exposure of the skin.

**Figure 4:**
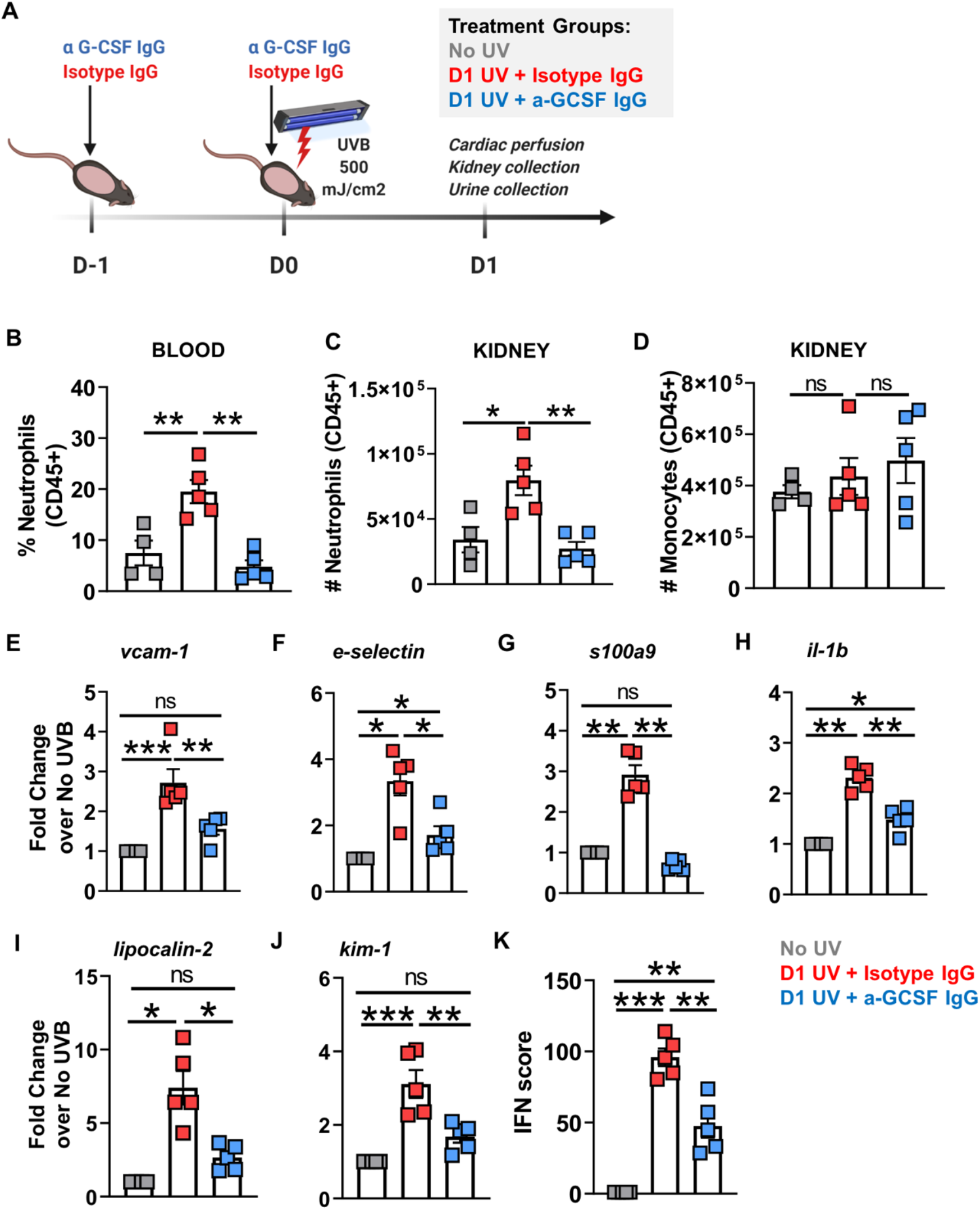
UVB light-triggered neutrophil migration to the kidney is required for kidney inflammatory and injury response. **(A)** B6 mice (n=4-5 per group) were either not exposed to UV (No UV, grey), or were exposed to UV and received 50 ug isotype control IgG i.p. (red), or were exposed to UV and received 50 ug anti-GSF IgG i.p (blue). Two doses of anti-G-CSF blocking IgG or isotype control IgG were given at 24h and 3h prior to UVB skin exposure. One day after UV exposure (D1), kidneys were collected and analyzed by flow cytometry and QPCR. (**B-D**) Flow cytometry analysis of (B) % neutrophils (CD11b+Ly6C+Ly6G+) in the blood, (C) number of neutrophils per kidney, and (D) number of monocytes (CD11b+Ly6C+Ly6G-) in the CD45+ kidney cell population. (**E-J**) Gene expression analysis of (E-F) adhesion molecules, *vcam-1* and *e-selectin*, (G-H) inflammatory mediators, *s100a9* and *il1b*, and (I-J) kidney injury markers, *lipocalin-2* and *kim-1*. (**K**) Type I interferon (IFN-I) score in the kidneys was evaluated based on relative expression of 10 interferon stimulated genes (ISGs) as described in Methods. (B-K) Statistical significance was determined by one-way ANOVA with multiple comparison and Bonferroni post-hoc (n=4-5 per group; *p < 0.05, **p < 0.01, ***p < 0.001, ns = not significant).

### A subpopulation of kidney neutrophils has the phenotype of reverse transmigrated cells, a phenotype also detected in the blood of SLE patients

While it has long been assumed that neutrophils entering sites of infection or sterile inflammation subsequently undergo apoptosis and are rapidly removed by macrophages^48,49^, a number of studies have revealed that some neutrophils traffic in a bi-directional manner, i.e. after entering a tissue, they transmigrate back into the bloodstream and infiltrate another location, a process referred to as reverse transmigration (rTM)^50-54^. To investigate whether neutrophils that homed to the kidney after skin exposure to UVB light transited via the exposed skin, we studied neutrophil migration in a photoactivatable B6 mouse model (UBC-PA-GFP^55^). The experimental scheme is illustrated in **Fig. 5A**: one day after skin exposure to a single dose of UVB light, the skin was exposed to violet light (405 nm) such that infiltrating immune cells were photoconverted to GFP-positive cells. Photoconverted neutrophils (GFP+Ly6G+) were then quantified in perfused kidneys the next day. Flow cytometry analysis revealed that ~20% of kidney infiltrating neutrophils were GFP-positive (**Fig. 5B**). In contrast to the models of sterile inflammation in the liver or cremaster muscle^50,53^, we did not detect photoconverted neutrophils in the lung (not shown), possibly because few neutrophils were found in the lung on day 2 (Fig. 1F).

**Figure 5:**
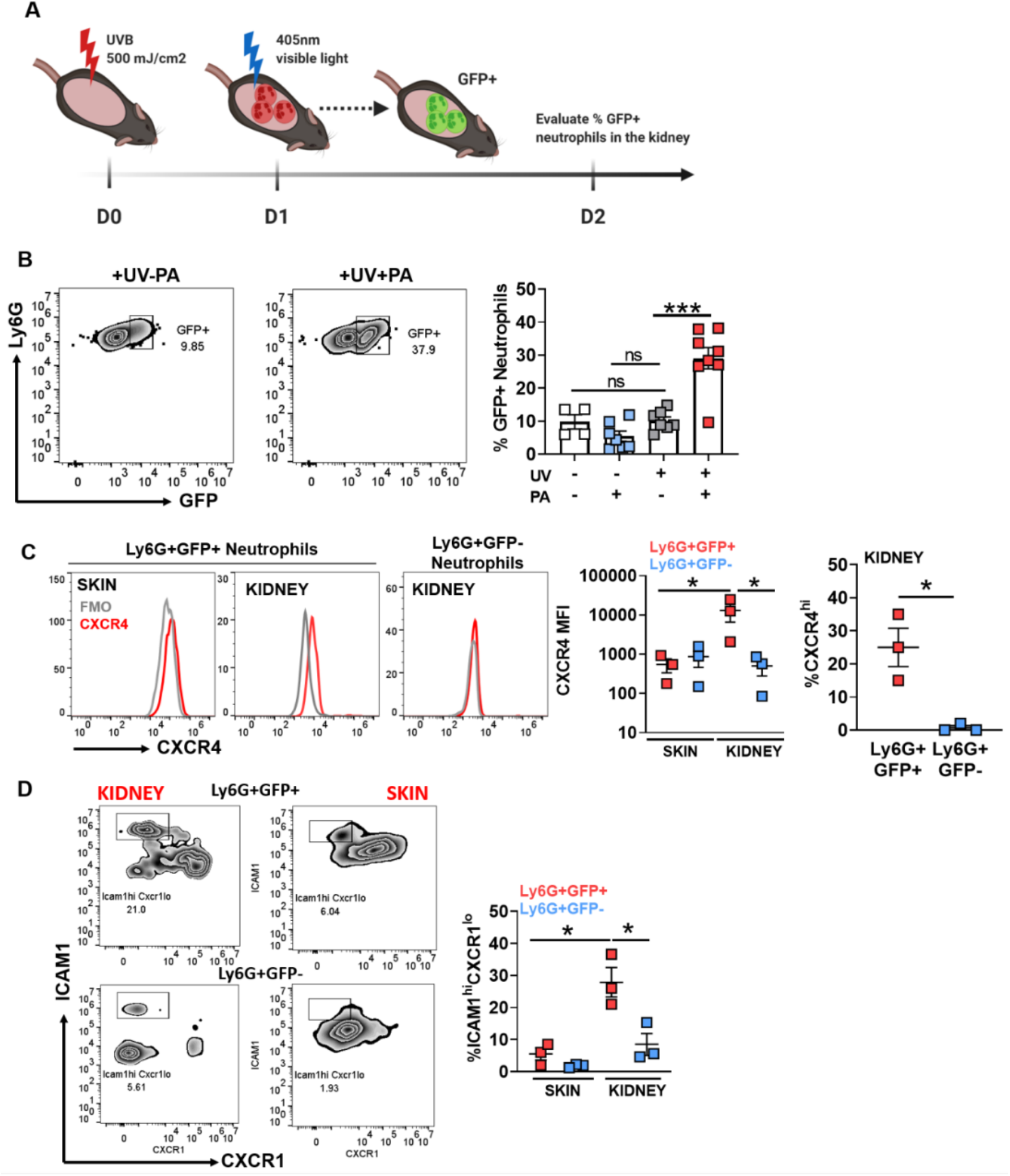
A subpopulation of kidney neutrophils are derived from UV light-exposed skin. **(A)** UBC-PA-GFP mice were shaved and irradiated with UVB light as in Fig.1. One day (D1) after UV exposure, the UV-irradiated skin was subjected to photoactivation (PA) by violet light (405 nm) as described in Methods. Twenty-four hours after PA (D2), cardiac perfusion was performed and kidneys collected. (**B**) The percentage of GFP+ neutrophils (CD45+Ly6C^int^Ly6G^hi^) in the kidneys of UV exposed (UV +) and photoactivated (PA +) mice was determined by flow cytometry (FC). The percentage of GFP+ neutrophils was compared to the kidneys from three different mouse controls: no UV, no PA; no UV, + PA; + UV, no PA by one-way ANOVA with multiple comparison and Bonferroni post-hoc (n = 4-8 animals/treatment, ***p < 0.001, ns = not significant). (**C**) Mean fluorescence intensity (MFI) and % CXCR4+ cells were determined by FC in photoconverted (GFP+, red) neutrophils in the skin and kidney and GFP-(blue) neutrophils in the skin and kidney of mice exposed to UV light and photoactivated as shown in panel A. (**D**) FC analysis of the % ICAM1^hi^CXCR1^lo^ cells in the GFP+ and GFP-neutrophil populations in the skin and kidneys of mice exposed to UV light and photoactivated as shown in panel A. (**C-D**) Statistical differences between groups were determined by Student’s t-test (n=3 individual experiments; *p < 0.05).

In previous studies, neutrophils having undertaken rTM were characterized by expression of surface markers such as CXCR4^hi^, also marking aged neutrophils^50,56-58^, and ICAM1^hi^CXCR1^lo^ neutrophils^52,59^. Indeed, GFP+ (rTM) kidney neutrophils expressed CXCR4, in contrast to low CXCR4 levels on GFP+ neutrophils in the skin or GFP-neutrophils in the kidney (**Fig. 5C**). Renal expression of CXCR4 ligand, *cxcl12*, 1-2 days after skin exposure to UVB light was concurrent with the presence of CXCR4^hi^ neutrophils in the kidney (Fig. 3F), possibly providing the chemotactic stimulus for recruitment of these neutrophils into the kidney^60,61^. Similarly, we found significantly greater levels of ICAM1^hi^CXCR1^lo^ GFP+ kidney neutrophils, compared to GFP+ skin and GFP-kidney neutrophils (**Fig. 5D**). Together, these data demonstrate that a subpopulation of neutrophils first enter the UV exposed skin, and then travel to the kidney. Some of these neutrophils express markers of aged and rTM neutrophils, which are associated with tissue injury at distal sites^50-53^ and reduced apoptosis^59^, and thus could contribute to the kidney inflammation resulting from acute skin exposure to UV light (Fig. 3). We also detected CXCR4^hi^ and ICAM1^hi^CXCR1^lo^ neutrophils in the bone marrow late after UV exposure (day 6, Supplementary Fig. 7), suggesting that these neutrophils home back to the bone marrow as previously reported in liver injury and virus infection of the skin^62,63^.

While both the aged (CXCR4^hi^) and the reverse migrating (ICAM1^hi^CXCR1^lo^) neutrophil phenotypes have been associated with inflammatory functions and tissue injury in different murine models^51,52,60^, few studies have been performed on these cell populations in human disease. Given the emerging role of neutrophils in SLE^11,15-17^ and their apparent heterogeneity^18^, we investigated whether PMNs or low density granulocytes (LDGs)^64,65^ from SLE patients demonstrated the phenotypes of aged, CXCR4^hi^, or reverse migrating, ICAM1^hi^CXCR1^lo^, neutrophils. Flow cytometry profiling of PMNs and LDGs revealed higher CXCR4 expression on the surface of these cells in SLE patients, compared to healthy controls (**Fig. 6A-C**). Moreover, we found a small, but significantly higher, percentage of ICAM1^hi^CXCR1^lo^ PMNs in SLE blood (mean 3.5%) relative to healthy controls (mean 1.4%) (**Fig. 6B**). Interestingly, the LDG neutrophil population (CD15^+^CD14^lo^CD10^+^ in PBMCs) contained a greater percentage of ICAM1^hi^CXCR1^lo^ cells, than the PMNs, in both healthy (mean 7.1%) and SLE blood (mean 29.4%) (Fig. 6B-D). The rTM-like population was significantly more highly represented in LDGs from SLE patients relative to healthy controls (**Fig. 6D**). These data are the first to identify the presence of aged and rTM neutrophil phenotypes in PMNs and particularly LDGs in SLE patients.

**Figure 6.**
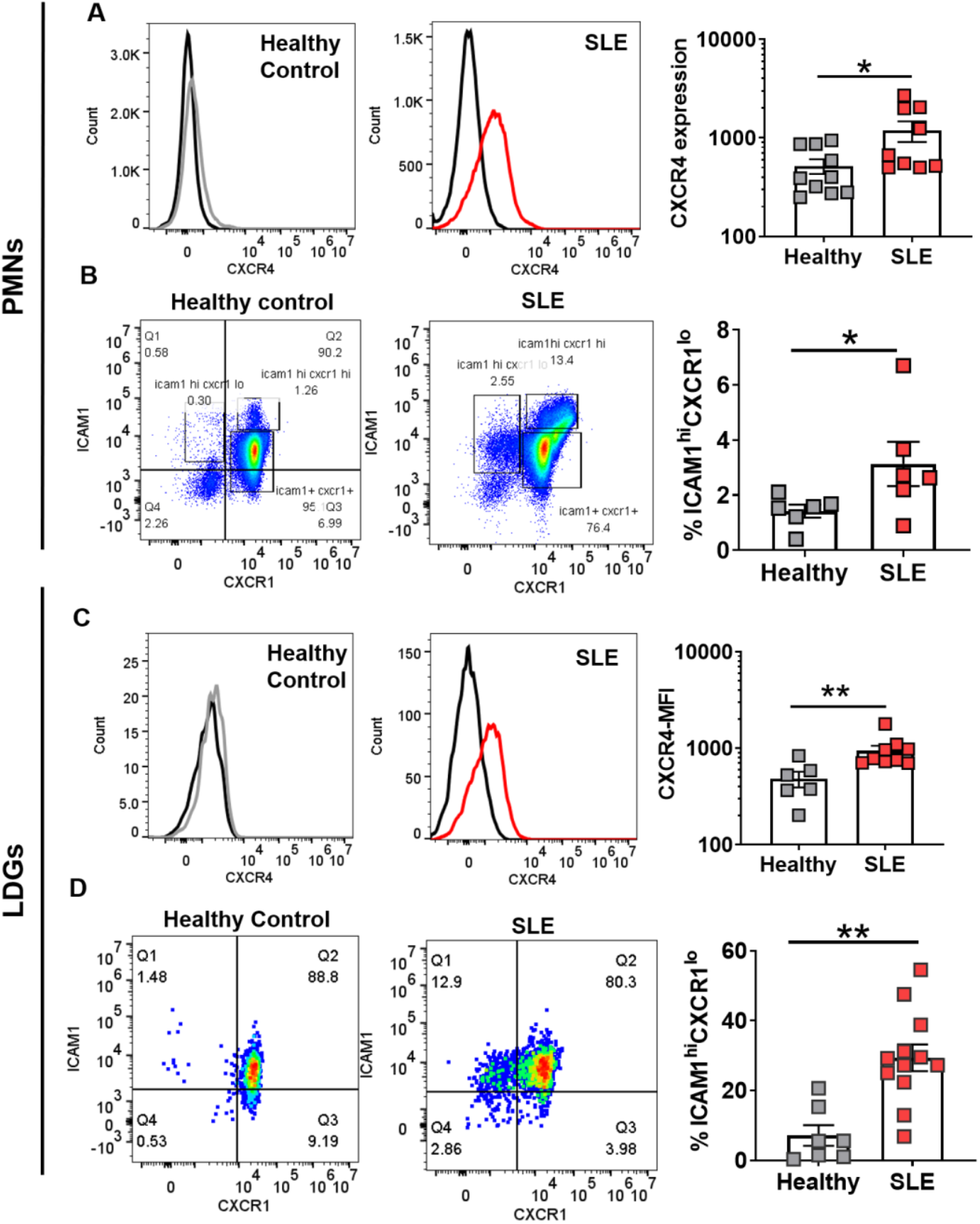
Increased CXCR4 expression and presence of ICAM1^hi^CXCR1^b^ neutrophils (PMNs) and low-density granulocytes (LDGs) in SLE patients. **(A, C)** Mean fluorescence intensity (MFI) of CXCR4 was determined by flow cytometry analysis of (**A**) circulating PMNs (CD66b+) and (**C**) Low density granulocytes (LDGs, CD10+CD15+CD14-) from healthy controls (grey) and SLE patients (red). (**B, D**) The percentage of ICAM1^hi^CXCR1^lo^ cells was determined by FC of (**B**) circulating PMNs and (**D**) LDGs from healthy controls and SLE patients. Representative histograms and gates are shown. Statistical differences between groups were determined by Student’s t-test (n=6-12; *p < 0.05, **p < 0.01).

## Discussion

Photosensitivity and its effects on systemic disease flares^4-6,66,67^ in SLE remain poorly understood. Here, we report several novel findings that could help explain how exposure to sunlight impacts distant organs such as the kidney. UVB light-mediated skin exposure mobilized neutrophils from the bone marrow that not only infiltrated the site of skin injury, but also accumulated in the kidneys. The neutrophil response was dependent on IL-17A and G-CSF. Neutrophils in the kidney had pro-inflammatory phenotypes as evidenced by increased expression of ROS, extracellular localization of neutrophil elastase, a higher proportion of ‘aged’ (CXCR4^hi^) as well as rTM (ICAM1^hi^CXCR1^lo^) cells. Neutrophils could be directly implicated in subclinical kidney injury since neutralization of G-CSF resulted in significant reductions in the expression of adhesion molecules, inflammatory cytokines, and kidney injury markers.

Other types of skin injury such as tape stripping and topical application of a TLR7 agonist have been reported to enhance kidney injury in certain lupus models, although disease enhancement was thought to be mediated by macrophages and dendritic cells rather than neutrophils^68,69^. Having demonstrated that tape stripping did not lead to neutrophil recruitment to the kidney, we propose that UV light exposure is different from other types of sterile skin inflammation and is more relevant to SLE. We found that exposure to UV light triggered a rapid and strong induction of *IL17a* in the skin together with high circulating IL17A protein levels (~100-fold increase), which was required for both local and systemic neutrophil migration after exposure to UV light. The concurrent 100-1000-fold induction in the skin *g-csf* expression suggests that the IL17-A/G-CSF axis is the likely mechanism responsible for neutrophilia and neutrophil tissue homing in response to UV light. The role of IL17A in regulating granulopoiesis and neutrophil recruitment via induction of G-CSF is well recognized under homeostatic conditions^34,70^ and a few studies have identified IL17-A as important for neutrophil recruitment to the sites of sterile inflammation^71,72^. In lupus, elevated IL-17A expression is found in cutaneous lesions^73,74^ and increased circulating IL-17A levels have been associated with worse disease manifestations^75,76^, although the specific relationship to neutrophils, a pathogenic population in this disease,^11,15-17^ remains unexplored. Besides its direct effects on G-CSF-mediated mobilization of neutrophils from the bone marrow, IL-17A may also contribute to neutrophil homing to the kidney by stimulating expression of adhesion molecules (e.g. VCAM-1 and E-Selectin) on the renal endothelium^77,78^. While under steady state conditions, phagocytosis of neutrophils downregulates IL-23 and subsequently IL-17-triggered neutrophil recruitment,^70^ the upstream pathways that regulate IL-17A expression in response to UV light are yet to be defined. Using the same model of acute UV exposure, we recently reported that a single dose of UV light triggered a local and systemic IFN-I response, which was required for efficient neutrophil recruitment to the skin^9^. Since IFN-I has been shown to induce IL-17A production^79^, it is plausible that these pathways either independently or in concert lead to UV-light mediated neutrophil migration to the skin we report here.

Neutrophils play multiple roles during tissue inflammation. When activated by PAMPS and DAMPS or by receptor engagement, neutrophils directly contribute to tissue injury by releasing proteases, reactive oxygen species (ROS), and extrusion of neutrophil extracellular traps (NETs)^80,81^. Neutrophils can also promote tissue repair by phagocytosis of cellular debris and by enabling tissue revascularization^50^. In SLE, a neutrophil gene signature is a strong predictor of active disease including LN and cutaneous lupus^15-17^. Neutrophils are found in cutaneous lesions^10-12^ as well as kidney biopsy specimens^13,14^ of SLE patients and their inflammatory properties have been attributed to NET formation^14,21^. This process includes increased production of ROS^19,82^, mitochondrial DNA release^19^, as well as release and induction of inflammatory proteins such as s100A9^83^, IL-1b^84^, and type I interferons^19,85,86^. Our findings that kidney infiltrating neutrophils produced ROS and that inhibiting neutrophil migration to the kidney abrogated s100A9 and significantly reduced IL-1b expression as well as the kidney IFN score, suggest that similar inflammatory functions might be at play. Relevant to kidney disease in SLE, neutrophils mainly localized to the tubulointerstitial (TI) endothelium, the site of increased expression of kidney injury markers *kim-1* and *lipocalin-2* in LN kidney tissues^87-89^. Since elevated Kim-1 and Lipocalin-2 are recognized markers of TI injury in SLE^40,42,90^, and blocking neutrophil migration to the kidney inhibited their expression, we can speculate that the transient proteinuria observed after UV exposure could be a consequence of neutrophil-mediated sub-clinical TI inflammation^90-92^. Increased kidney *vcam-1* expression, another marker of TI injury in SLE^93^, further supports this model. Whether subclinical glomerular damage, resulting in increased *cxcl12* expression in the kidney (discussed below)^94^, might contribute to the observed neutrophil recruitment and proteinuria remains to be determined.

In addition to propagating inflammation at the local sites of sterile or infectious injury, neutrophils have been shown to home to distant organs^50,53,95,96^ where, depending on the context, they contribute to lung tissue injury via ROS production^52^ or activate the adaptive immune system in the lymph nodes and the bone marrow^63,95^. The inflammatory nature of these neutrophils has been attributed to their ability to exit the primary site of injury and migrate to secondary sites, i.e. to undergo reverse transmigration (rTM)^51-54,95^. Our findings that photoactivation of GFP containing cells in the skin following UV exposure led to detection of GFP+ neutrophils in the kidney with the phenotype of rTM, suggest that a subpopulation of renal neutrophils reverse migrated from UV-exposed skin. rTM have in other circumstances been shown to be pro-inflammatory^52,53^ and have reduced apoptosis^59^, leading to the possibility that they play a role in UV-induced kidney injury. The elevated CXCR4 expression on these neutrophils is particularly relevant as increased renal expression of *cxcl12*, a CXCR4 ligand expressed by podocytes and tubules^94,97^, was observed, likely providing a chemotactic signal for this neutrophil population. Increased CXCR4 expression on immune cells coupled with high levels of renal CXCL12 have been identified in SLE patients with LN^98^ and this pathway was implicated in lupus as well as ischemia reperfusion kidney injury in mice^94,99,100^.

Tissue specific neutrophil migration and the mechanisms responsible for differential expression of adhesion molecules in different organs is not well understood^101,102^. Higher levels of expression of the chemokine *cxcl12* in the kidney, but not the lung, could explain the preferential presence of CXCR4^hi^ neutrophils in the kidney. In addition, the induction of expression of *vcam-1* and *e-selectin* in the kidney, compared to no change in expression in the lung, likely contributes to preferential neutrophil recruitment and retention in the kidney. The reduction in neutrophil infiltration of the kidney following neutralization of G-CSF can be attributed to both a reduction in circulating numbers and activation of neutrophils as well as a reduction in local expression of *vcam*-1 and *e-selectin*. However, we cannot be certain whether initial upregulation of these adhesion molecules following skin UV exposure is attributed to skin-derived cytokines /DAMPS or whether neutrophils, through release of serine proteases, contribute directly to their expression as has been shown in other contexts.^103^ Regardless of the mechanisms, and noting that we have not performed a comprehensive analysis of all tissues, our studies suggest organ selectivity in UV-light mediated systemic effects.

In patients with SLE, it is difficult to unequivocally relate skin UV light exposure to exacerbations of systemic disease as there is significant variation (1-3 weeks) in visible photosensitivity responses in different individuals^28,104^. In addition, subclinical kidney injury may readily be missed until consecutive exposures to UV light compound the effects. While we found that neutrophil-mediated kidney inflammation in response to UV light does not cause clinical disease in healthy mice, such a mechanism may contribute to LN flares in photosensitive lupus patients in multiple ways: Fc receptor engagement by immune complexes could enhance neutrophil recruitment resulting in ROS and protease release^105,106^; the heightened capacity of lupus neutrophils and LDGs to produce NETs, which in SLE patients are not cleared efficiently^107,108^, could lead to release of tissue-damaging proteases^109,110^, propagation of the IFNI response^85,86^, or direct damage to the kidney endothelium by creating vascular damage and leakage^14,111^. Moreover, the underlying differences in lupus skin, such as enhanced IFN-I signaling^7,112,113^ and defects in protective Langerhans cell population^114^ could inform the extent and nature of neutrophil-mediated systemic responses. The exact mechanism might in addition be influenced by the neutrophil/LDG phenotype, as heterogeneity within these populations has become more apparent in SLE^65^. Our findings of elevated CXCR4 and ICAM1^hi^CXCR1^lo^ (rTM) circulating populations, particularly in SLE LDGs, further add to this heterogeneity and suggest that some of the more pro-inflammatory granulocytes in blood might have prior “tissue-experience”, i.e. residence in the affected organ tissues (e.g. skin) prior to systemic dissemination. These phenotypes have previously been reported in synovial fluid and circulating neutrophils of rheumatoid (RA) patients^59,115^ and in acute pancreatitis-associated lung injury^116^. Their role in SLE warrants further investigation.

## Materials and Methods

### Mice

All animal experiments were approved by the Institutional Animal Care and Use Committee of the University of Washington, Seattle. Initial experiments were conducted using both male and female C57BL/6J mice (Jackson Labs, 3-4 months old, Figs. 1-2). Female mice were used for all subsequent experiments. Photoactivation studies were performed on B6.Cg-*Ptprc^a^* Tg(UBC-PA-GFP)1Mnz/J female mice (Jackson Labs, 3-4 months old) ^55^. The animals were housed in pathogen-free conditions and maintained in light-dark cycles of 12 hrs with ad libitum access to food and water. The experiments on all the animals were performed at the same time of the day, as to avoid the influence of circadian rhythm on neutrophil migration ^56^.

### Exposure to UVB light and tape stripping

The dorsal aspect of male and female C57BL/6 mice was shaved at least 24hr prior to irradiation with UVB light and only the mice with non-pigmented skin were used in the experiments. Mice were anesthetized with isoflurane and exposed to one dose of UVB light (500 mJ/cm2) using FS40T12/UVB bulbs (National Biological Corporation), with peak emission between 300 and 315 nm. The UVB light energy at the dorsal surface was measured with Photolight IL1400A radiometer equipped with a SEL240/UVB detector (International Light Technologies). Mice were euthanized one (D1), two (D2), or six (D6) days after exposure to UVB light and non-irradiated mice very used as controls (D-1) (Figure 1A). Epidermal tape stripping was performed as previously described^22^ and skin evaluated 24hr after injury.

### Inhibition of IL-17A and G-CSF

Mice were treated i.v. with 100 μg purified rat anti-mouse IL-17A IgG (Biolegend) or isotype control 3hr before exposure to UV light (1 x 500 mJ/cm^2^) (Fig. 2I). Neutrophil presence in the blood, skin, and saline-perfused kidney tissue was evaluated by flow cytometry 24hr after UV exposure as described below. Neutrophil migration in response to UV light was inhibited by treating mice intraperitoneally (i.p.) with 50 μg anti-mouse G-CSF monoclonal antibody or mouse IgG isotype control (R&D Systems) 24h and 3h prior to skin exposure to UVB light (1 x 500 mJ/cm^2^) (Fig. 4A). Following cardiac perfusion, neutrophil and monocyte numbers in the kidney were evaluated by flow cytometry and gene expression by qPCR, as described below. Mice not exposed to UVB light were used as an additional control group.

### Photoactivation studies

UBC-PA-GFP mice were shaved as above and a portion of the back was irradiated with one dose UVB light (500 mJ/cm^2^) as above. Only the skin area that will be photoconverted was exposed to UVB light, while the remaining skin was covered with aluminum foil. Twenty-four hours after UVB-induced sterile injury, the entirety of the UVB-irradiated skin region was subjected to violet light (405 nm) using 50W LED lamp with 0.37NA fiber (Mightex) at the power of 100 mW. The method used for photoactivation (PA) was optimized based on a published protocol of skin-specific photoconversion ^117^ Twenty-four hours after PA, kidneys and lung were collected and processed for flow cytometry analysis as described below. The approach is outlined in the diagram in Figure 5A. Percent GFP+ neutrophils in the kidney was compared to those found in mice under three control conditions: i) no UV and no PA, ii) UV but no PA, and iii) no UV but PA.

### Isolation of cells from different organs

Following euthanasia with CO2 inhalation, a piece of dorsal skin was removed and kept in RPMI on ice until processing. Blood was collected by cardiac puncture in EDTA coated syringes. Cardiac perfusion was subsequently performed using phosphate buffered saline (PBS) (~60ml) through the left ventricle after snipping the right atrium. A successful perfusion was determined by observing complete kidney and lung pallor. Lungs, kidneys, spleen, and both femurs were removed carefully and placed in RPMI on ice until sample processing. Cells were isolated from equivalent areas of skin tissue (~30mm^2^) by mincing and digesting the tissue with 0.28 units/ml Liberase TM (Roche) and 0.1 mg/ml of Deoxyribonuclease I (Worthington) in PBS with Ca^+2^ and Mg^+2^ for 60 minutes at 37°C with shaking. Cells were isolated from one kidney per animal; kidney tissues were minced and digested in 2 mg/ml collagenase type I (Worthington) for 30 min at 37 °C according to a 118 +2 +2 published protocol^118^. Lung cells were harvested by digesting tissue in PBS (Ca^+2^ and Mg^+2^) with 1 mg/mL collagenase type 1 and 60 U Deoxyribonuclease I as described ^119^. Whole spleens were crushed onto a 40 μm cell strainer and washed with RPMI. Bone marrow cells were isolated from femurs as previously described ^120^. Whole blood and cells from the spleen, bone marrow, kidneys, and lungs were treated with 1X red blood cells lysis buffer (BD Biosciences). Following isolation/digestion, cells from all tissues were filtered (40 μm), washed with PBS, and resuspended in RPMI prior to counting.

### Flow cytometry staining and analysis

Cells were treated with Fc Block TruStain FcX (Biolegend) and stained with Zombie Aqua viability dye (Biolegend) per vendor’s recommendations. Cells were washed and surface staining performed using mouse–specific fluorescent antibodies purchased from Biolegend. Different myeloid cell populations were analyzed in the live immune cells (Zombie Aqua-CD45+) gate. Percent and number of neutrophils (Ly6C^int^Ly6G^hi^) and monocytes (Ly6C+Ly6G-) were quantified. Representative flow cytometry gating is presented in Supplementary Fig. 8. Percent expression and mean fluorescence intensities (MFI) of surface markers in the photoactivatable model (CXCR4, ICAM1, and CXCR1) were determined using fluorescence minus one (FMO) staining controls in the neutrophil gate (Ly6G^hi^). Production of reactive oxygen species (ROS) in kidney neutrophils was evaluated *ex vivo* with 1,2,3 dyhidroxyrhodamine (DHR) dye (Sigma Aldrich). Single cells isolated from kidneys of mice not exposed to UV light (D-1) or one day after UV exposure (D1) were incubated with 5 μM DHR in RPMI for 20 minutes at 37C. Mean fluorescence intensity of DHR was evaluated by flow cytometry, relative to MFI of cells incubated without DHR. All samples were processed using CytoFLEX flow cytometer (Beckman Coulter) and data analyzed with FlowJo software v10 (Tree Star).

### Gene expression and proteomic analyses

Skin, kidney, and lung samples were stored in RNAse Later solution (Qiagen). RNA was extracted by RNA Easy kit from Qiagen (Valencia, CA) and cDNA was synthesized using High Capacity cDNA synthesis kit (Applied Biosciences). The transcripts of inflammatory chemokines, cytokines, and adhesion molecules were quantified by real time quantitative PCR using the primers listed in Supplementary Table 1 and normalized to the average of *18s* transcript levels. The dose of UVB light used did not affect *18s* expression in any of the organs. Relative expression of mRNA targets was determined using the standard formula, 2^(-Ct) x coefficient. IFN scores were derived from expression of 10 representative interferon stimulated genes (ISG, *Mx1, Irf7, Isg15, Isg20, Ifi44, ifit1, ifit3, oasl1, usp18*, and *ifi27l2a*) as previously described ^121^. The mean and standard deviation (SD) level of each ISG were determined in the kidneys of mice not exposed to UV light (mean_noUV_ and SD_noUV_). These were then used to standardize the expression levels of each ISG in the kidneys of treated mice (IgG isotype + UVB or a-GCSF + UVB). The standardized expression levels were then summed for each mouse to derive an IFN expression score; *i* = expression of each ISG; Gene i_treatment_= relative gene expression level after treatment; Gene i_noUV_ = relative gene expression level in mice not exposed to UVB light; SD = standard deviation. 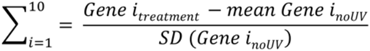 Protein levels in plasma and skin tissue protein extracts were measured using defined analyte panels: Legendplex Mouse Inflammation Panel and Inflammatory Chemokine Panel (Biolegend).

### Immunofluorescence staining of frozen kidney tissues

Saline-perfused kidneys were fixed in 4% paraformaldehyde for 1hr at RT and, following immersion in 30% sucrose, frozen in optimal cutting medium (OCT, Sakura Finetek). 5 μm tissue sections were rehydrated in PBS prior to immunofluorescence (IF) staining. Tissues were permeabilized with 0.1% Triton X-100 in PBS and subsequently blocked in PBS+0.1%Triton-X 100/5%BSA/5% rabbit serum. Neutrophil elastase (NE) and endothelial cells were detected by staining with anti-mouse NE Cy5 (1:100, Bioss) and anti-mouse PECAM-1/CD31 Al488 (1:100, Biolegend), respectively, at 4C overnight. Neutrophil presence in the kidney was confirmed by staining with anti-mouse Ly6G Al647 (1:100, Biolegend). Tissues were mounted using Prolong Gold with DAPI mounting medium (Invitrogen) and imaged with Nikon Eclipse 90i fluorescent microscope (Histology and Imaging Core, University of Washington).

### Skin histologic evaluation

Formalin fixed, paraffin embedded skin tissues sections (4-5μm) were stained with hematoxylin and eosin (HE) at the University of Washington Histology and Imaging Core at Day 0, Day 1, Day 2, and Day 6 following UVB irradiation (n=5 females). Skin was scored in a blinded fashion for the following parameters: inflammation of the dermis/subcutis; epidermal cell death; acanthosis; hyperkeratosis; serocellular crust formation; and erosion/ulceration. These parameters were scored on a scale of 0-4 depending on severity and extent with the exception of erosion/ulceration, which was scored as present (1) or absent (0). Representative images were taken using NIS-Elements BR 3.2 64-bit and plated in Adobe Photoshop with lighting adjustments applied to the entire image. Original magnification is stated.

### Urine measurements

Urine protein levels were evaluated by Bradford protein assay (ThermoFischer Scientific). Urine samples were tested at 1:40 dilution in PBS and protein concentration in urine determined based on serial dilutions of known concentrations of bovine serum albumin (BSA). Urine albumin/creatinine ratio (UACR) was evaluated by evaluated using Albuwell albumin assay and its companion Creatinine kit (Exocell) per manufacturer’s instructions.

### Human sample collection and flow cytometry analysis

Whole blood was collected in heparinized tubes from healthy volunteers and SLE patients following informed consent in the IRB-approved protocol (University of Washington; STUDY00001145). Neutrophils (polymorphonuclear cells, PMNs) and peripheral blood mononuclear cells (PBMCs) were isolated by ficoll paque discontinuous gradient separation (GE Healthcare). PMNs were purified from the erythrocyte pellet by 5% dextran sedimentation. Following RBC lysis, cells were stained with fluorescently labeled antibodies purchased from Biolegend. CXCR4 expression (aged phenotype, ^50,56^and percent ICAM1^hi^CXCR1^lo^ cells (rTM phenotype, ^59^) were evaluated in PMNs (CD66b+) and low density granulocytes [LDGs, CD15^+^, CD14^lo^, CD10^+^ in PBMCs; ^18^]. Samples were processed using CytoFLEX flow cytometer (Beckman Coulter) and data analyzed with FlowJo software v10 (Tree Star).

### Statistical analyses

Data were analyzed using GraphPad Prism 7 software (GraphPad Software Inc.) and presented as mean + SEM. Statistical difference between two data groups was determined relative to nonirradiated controls (D-1) using Student’s *t*-test. One way-ANOVA with multiple comparison and Bonferroni post-hoc was used to determine statistical significance between three groups in human UV exposure studies. Statistical significance was designated as follows: *p < 0.05, ** p < 0.01, ***p < 0.001.

## Acknowledgments

This work was funded by the National Institutes of Health Awards R56 AR073848 (K.B.E.), T32 AR007108 (S.SG.), R01 AR074939 (T.M.), and R21 AR075134 (T.M.); Human studies were funded by ITHS Catalyst Award UW (S.SG.). We thank Drs. Eric Butz, Jeffrey Ledbetter, Jessica Hamerman, Christian Lood, and Erica Noss for insightful discussion. We thank Drs. Stuart Shankland and Jonathan Himmelfarb for advice and discussion of kidney studies.

## Author contributions

S.SG. and K.B.E. conceived the study and wrote the manuscript. S.SG., L.T., X.S., J.T., and J.K. conducted the experiments. S.SG. and J.T. analyzed the data. T.M. and P.K. contributed to the conceptualization of the study and experimental design. J.S. performed pathology studies. All authors critically reviewed and edited the manuscript.

