## Supplementary Material for "Neutrophils Mediate Kidney Inflammation Following Acute Skin Exposure to UVB Light"

**Supplementary Figures:**

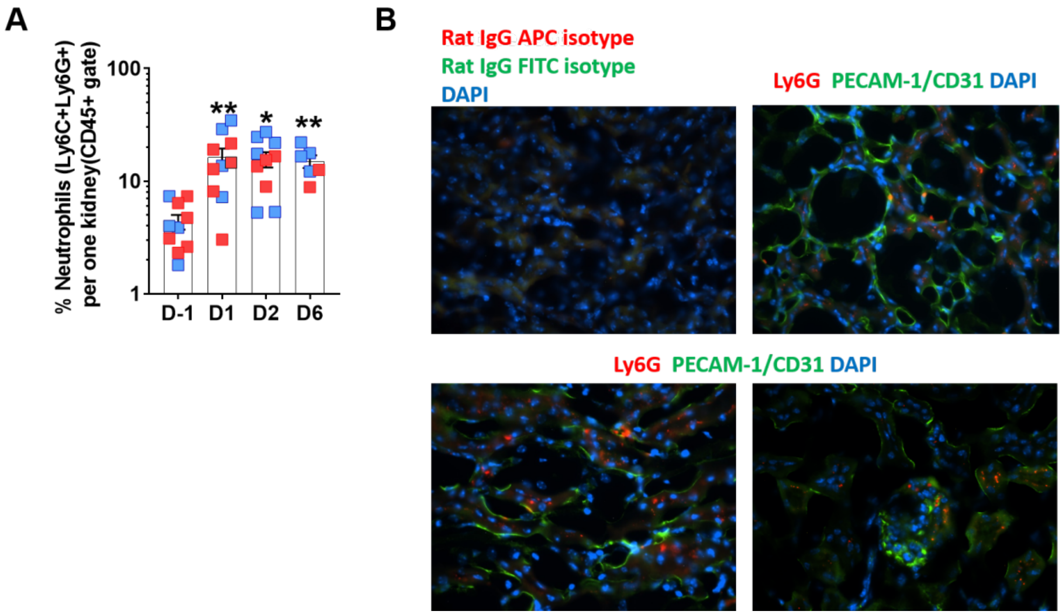


**Supplementary Fig. 1: (A)** Percent neutrophils in the CD45+ cells from perfused kidney tissue determined by flow cytometry prior to (D-1) and on different days (D1-6) after UV exposure as in Fig. 1, in female (red) and male (blue) mice. Statistical significance was determined relative to non-irradiated controls (D-1) using Student's t-test (n=6 -10 per group; *p<0.05, **p<0.01). **(B)** Evidence of kidney infiltrating neutrophils by immunofluorescene in kidneys on day 2 after UV exposure, demonstrating Ly6G (Cy5, magenta), PECAM-1/CD31 (Al488, green), and nuclear DAPI (blue) staining in the tubulointerstitium (top right, bottom left) and glomeruli (bottom right). Isotype control staining is shown in the top left panel. Magnification = 40X.

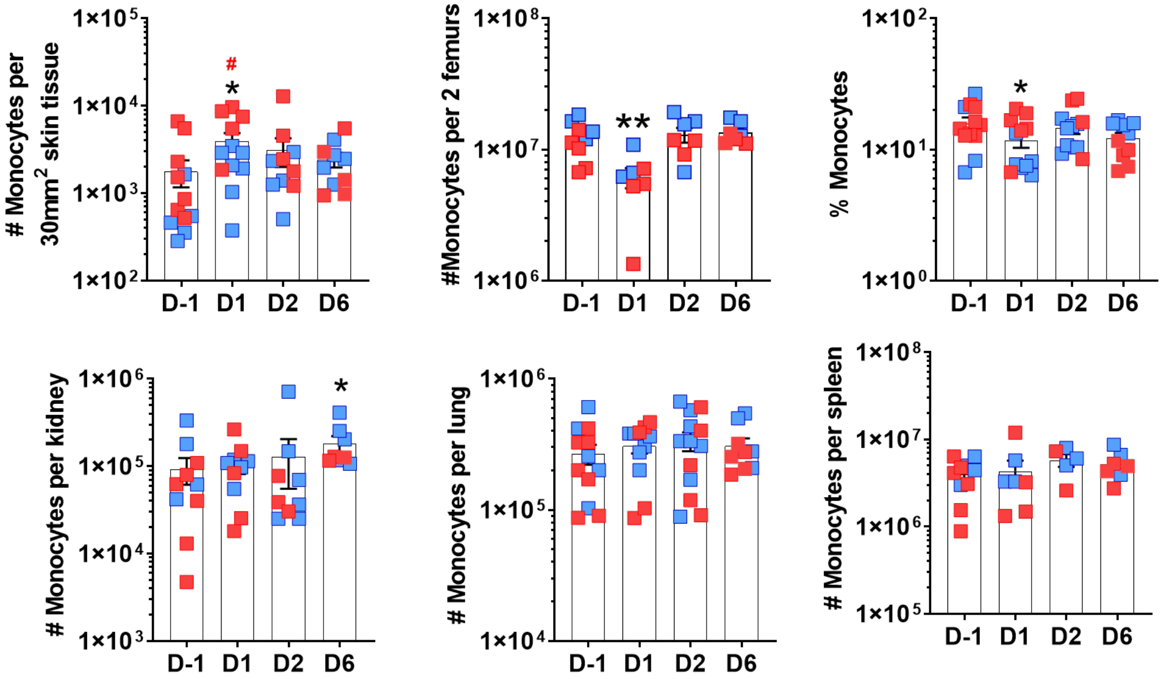


**Supplementary Fig. 2:** Number and percent monocytes in different organs determined by flow cytometry prior to (D-1) and on different days (D1-6) after UV exposure as in Fig. 1, in female (red) and male (blue) mice. Statistical significance was determined relative to non-irradiated controls (D-1) using Student's t-test (n=6 -10 per group; *p<0.05, **p<0.01).

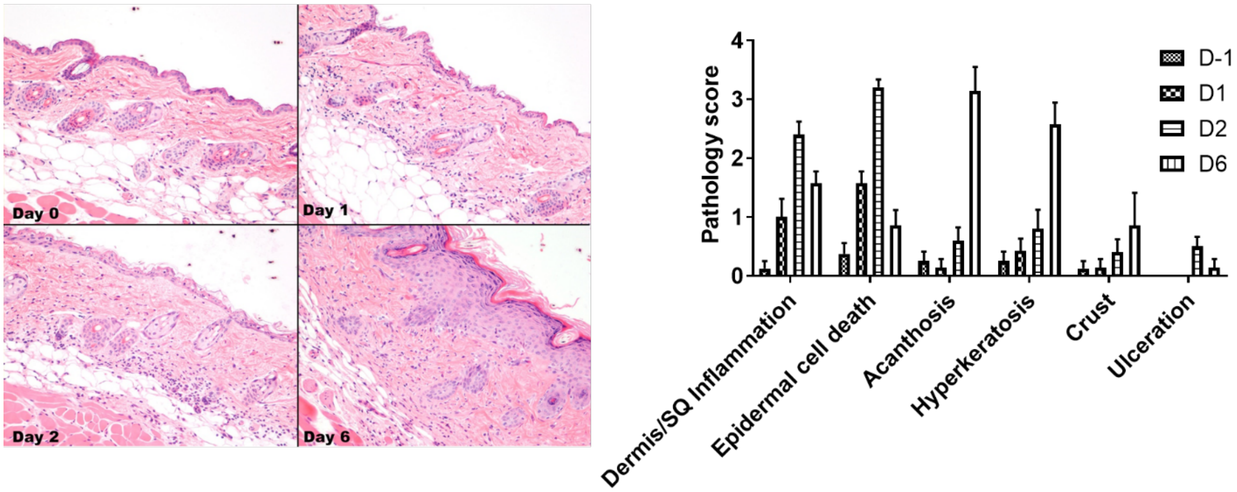


**Supplementary Fig. 3:** Representative skin pathology following H&E staining of non UV-exposed skin (Day 0) or skin taken on different days (1-6) after UV light exposure. Pathology scores quantified on the right (n = 5 per time point).

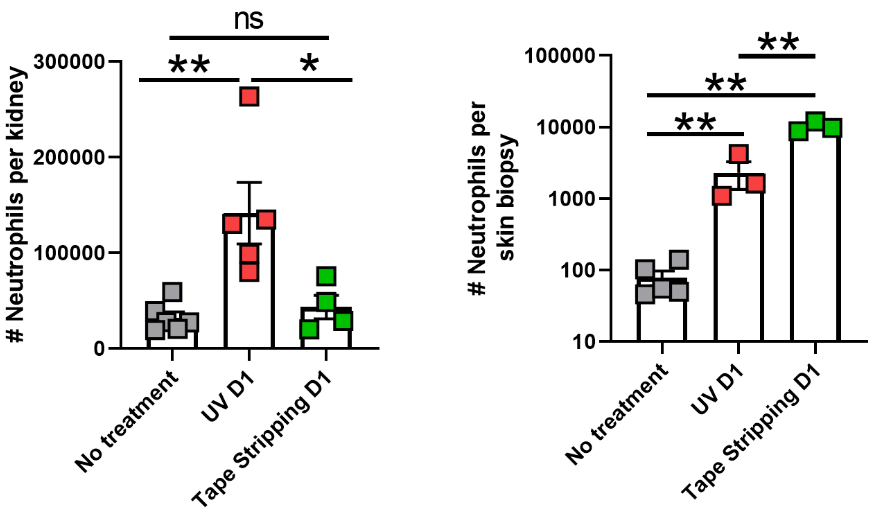


**Supplementary Fig. 4:** Number of neutrophil in kidney (left) and skin (right) were determined by flow cytometry (CD45+ live cell gate) in tissues from untreated mice, one day (D1) after UV exposure as in Fig. 1 or one day after epidermal tape stripping as described in the Methods. Statistical significance was determined by one-way ANOVA with multiple comparison and Bonferroni post-hoc (n=4 -6 per group; *$p<0.05$, **$p<0.01$, ns = not significant).

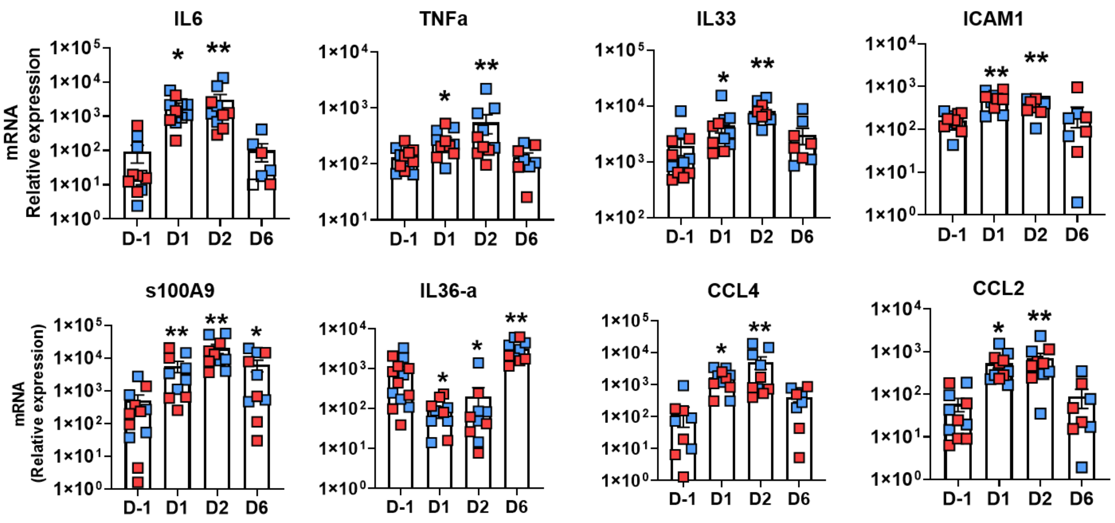


**Supplementary Fig. 5:** Relative mRNA expression in skin obtained from female (red) and male (blue) mice was quantified by QPCR using the primers listed in Table 1 and normalized to *18S* transcript levels. Bars represent mean relative expression ± SEM for all samples combined. Statistical significance was determined relative to non-irradiated controls (D-1) using Student's t-test (n=6 -10 per group; *p<0.05, **p<0.01).

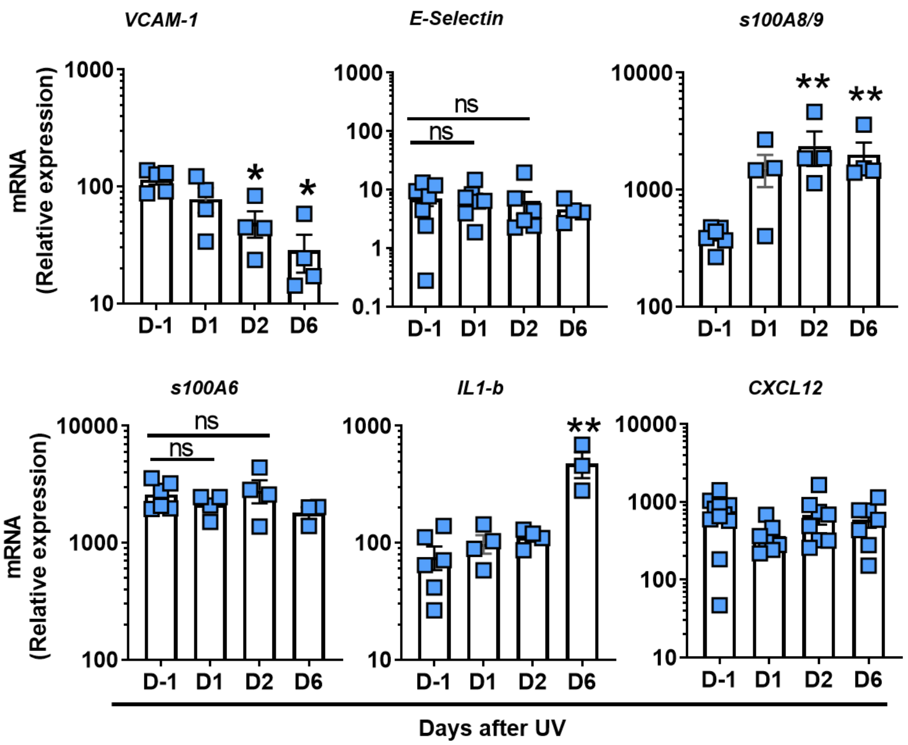


**Supplementary Fig. 6:** Relative mRNA expression in the perfused lung tissue prior to (D-1) or after skin exposure to UV light (D1-6) as in Fig.1. Statistical significance was determined relative to non-irradiated controls (D-1) using Student's t-test (n=3 -6 per group; *p<0.05, **p<0.01; ns = not significant).

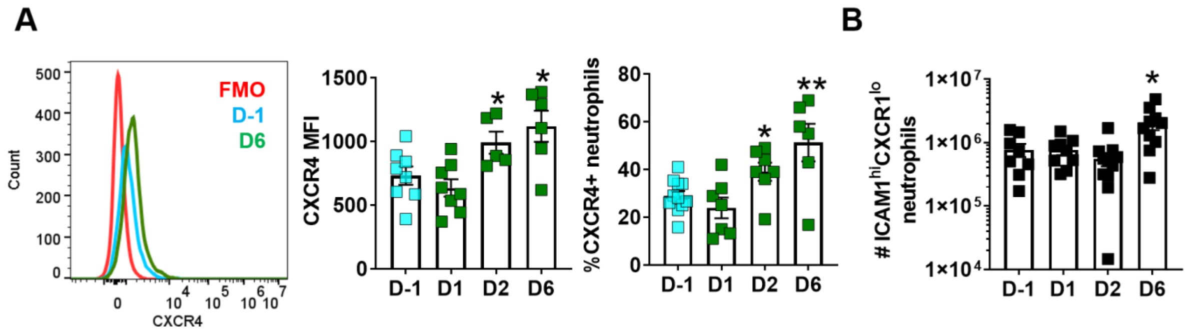


**Supplementary Fig. 7: (A)** CXCR4 mean fluorescence intensity (MFI) and % CXCR4 positive neutrophils in the bone marrow prior to (D-1) and on different days (D1-6) after skin exposure to UV light as in Fig. 1. Representative histograms in the leftmost panel. (**B**) Number of $ICAM1^{hi}CXCR4^{lo}$ neutrophils in the bone marrow prior to and after UV exposure.

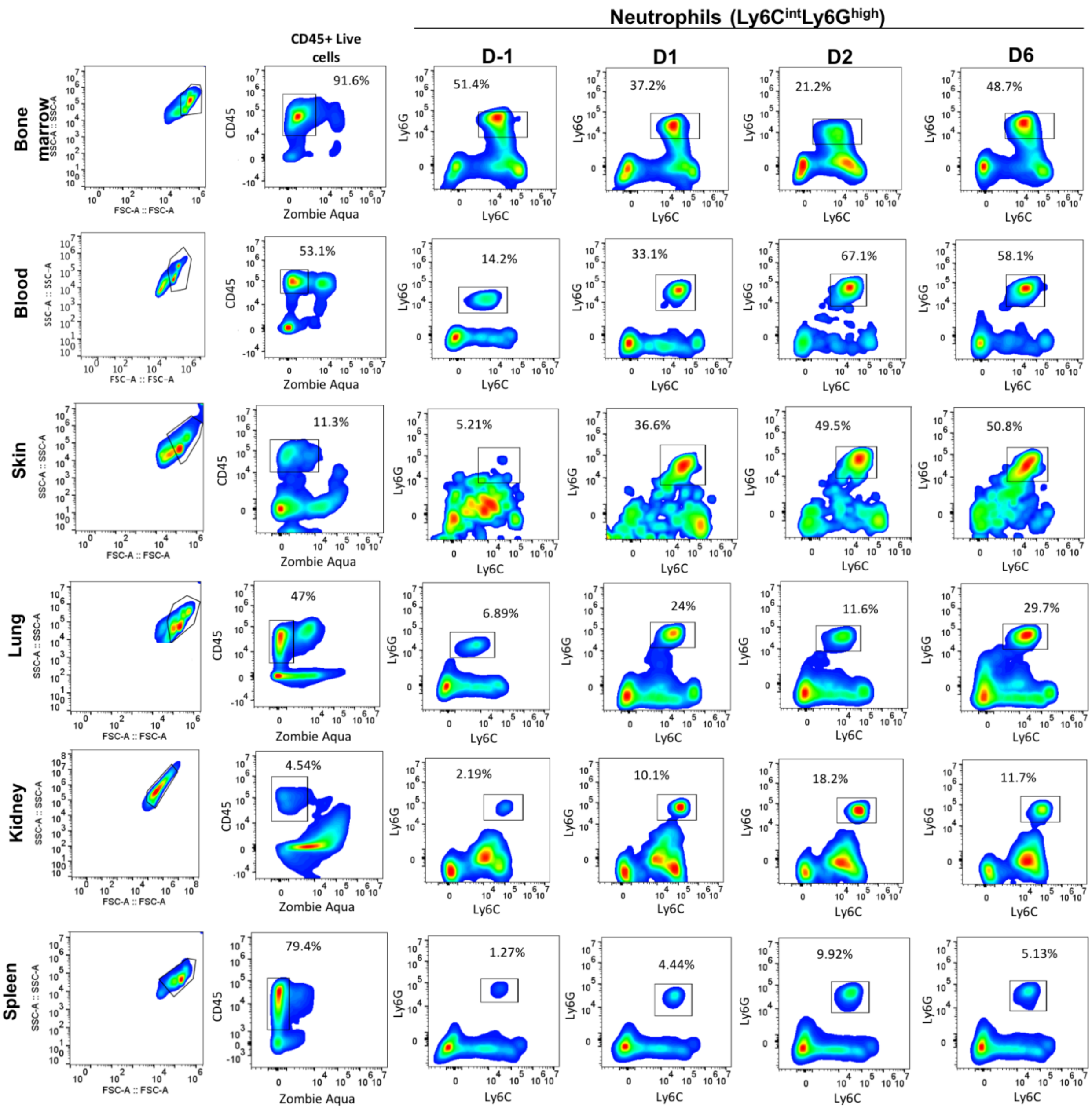


**Supplementary Fig. 8:** Representative flow cytometry gating of neutrophils (Ly6C$^{int}$Ly6G$^{high}$) in CD45+live cell gate in different tissues prior to (D-1) and on different days (D1-6) after skin exposure to UV light as in Fig. 1.

**Supplementary Table 1: Primers used for gene expression analysis**

| | Primer | Forward (5'--> 3') | Reverse (5'--> 3') |
|---|---|---|---|
| 1 | 18s | AACTTTCGATGGTAGTCGCCGT | TCCTTGGATGTGGTAGCCGTTT |
| 2 | TNF | CTGAACTTCGGGGTGATCGG | GGCTTGTCACTCGAATTTTGAGA |
| 3 | IL-6 | TCTATACCACTTCACAAGTCGGA | GAATTGCCATTGCACAACTCTTT |
| 4 | IL1b | TCCAGGATGAGGACATGAGCAC | GAACGTCACACACCAGCAGGTTA |
| 6 | IL33 | TCCAACTCCAAGATTTCCCCG | CATGCAGTAGACATGGCAGAA |
| 7 | IL36a | TGCCCACTCATTCTGACCCA | GTGCCACAGAGCAATGTGTC |
| 8 | G-CSF | TGCACTATGGTCAGGACGAG | GGGGTGACACAGCTTGTAGG |
| 9 | CCL4 | AACAACATGAAGCTCTGCGT | AGAAACAGCAGGAAGTGGGA |
| 10 | CXCL1 | CAATGAGCTGCGCTGTCAGTG | CTTGGGGACACCTTTTAGCATC |
| 11 | CXCL2 | CCAAGGGTTGACTTCAAGAAC | AGCGAGGCACATCAGGTACG |
| 12 | CXCL5/6 | TCCAGCTCGCCATTCATGC | TTGCGGCTATGACTGAGGAAG |
| 13 | CXCL12 | TGCATCAGTGACGGTAAACCA | CACAGTTTGGAGTGTTGAGGAT |
| 15 | ICAM-1 | TGTTTCCTGCCTCTGAAGC | CTTCGTTTGTGATCCTCCG |
| 16 | CCL2 | CCCAATGAGTAGGCTGGAGA | AAAATGGATCCACACCTTGC |
| 17 | S100A9 | CACAGTTGGCAACCTTTATG | CAGCTGATTGTCCTGGTTTG |
| 18 | S100A6 | TGAGCAAGAAGGAGCTGAAGGAGT | TTCTGATCCTTGTTACGGTCCAGA |
| 19 | VCAM1 | CCCAGGTGGAGGTCTACTCA | CAGGATTTTGGGAGCTGGTA |
| 20 | E-selectin | AGCTACCCATGGAACACGAC | ACGCAAGTTCTCCAGCTGTT |
| 21 | Lipocalin-2 | CAAGCAATACTTCAAAATTACCCTGTA | GCAAAGCGGGTGAAACGTT |
| 22 | Kim-1 | TGGCACTGTGACATCCTCAGA | GCAACGGACATGCCAACATA |
| 23 | Isg15 | AAGCAGCCAGAAGCAGACTC | CACCAATCTTCTGGGCAATC |
| 24 | Isg20 | TCACGGACTACAGAACCCAAG | TATCCTCCTTCAGGGCATTG |
| 25 | Ifit1 | TGCTGAGATGGACTGTGAGG | CTCCACTTTCAGAGCCTTCG |
| 26 | Ifit3 | TGGCCTACATAAAGCACCTAGATGG | CGCAAACTTTTGGCAAACTTGTCT |
| 27 | Irf7 | GTCTCGGCTTGTGCTTGTCT | CCAGGTCCATGAGGAAGTGT |
| 28 | Mx1 | CCTCAGGCTAGATGGCAAG | GGCAGACACCACATACAACC |
| 29 | Ifi44 | AACTGACTGCTCGCAATAATGT | GTAACACAGCAATGCCTCTTGT |
| 30 | Oasl1 | CAGGAGCTGTACGGCTTCC | CCTACCTTGAGTACCTTGAGCAC |
| 31 | Usp18 | TTGGGCTCCTGAGGAAACC | CGATGTTGTGTAAACCAACCAGA |
| 32 | Ifi27l2a | CTGTTTGGCTCTGCCATAGGAG | CCTAGGATGGCATTTGTTGATGTGG |
